# Human biting *Culex pipiens* bioform *molestus* is spread in several areas in south Sweden

**DOI:** 10.1101/616706

**Authors:** Luande Verah Nafula, Disa Eklöf, Anders Lindström, Steven Ger Nyanjom, Magnus Evander, Tobias Lilja

## Abstract

The mosquito species *Culex pipiens* is a potential vector of several pathogens infecting humans and occurs in two distinct bioforms, *pipiens* and *molestus*. Traditional morphological identification fails to separate the bioforms of *Cx. pipiens* despite their behavioural differences since they are morphologically indistinguishable. However, molecular methods can identify the two bioforms. The bioform *molestus* thrives in urban environments and bite all kinds of vertebrates, whereas bioform *pipiens* is more rural and mainly feed on birds.

Mosquito samples submitted in a citizen science project from people experiencing mosquito problems in South Sweden were analyzed to determine the geographical distribution of the *molestus* bioform of *Cx. pipiens*. Mosquito specimens were identified to species by DNA barcoding of the cytochrome C oxidase subunit I (COI) gene and the bioforms were determined with the CQ11 microsatellite marker. To establish other differences between the bioforms, part of the CPIJ001674 gene was sequenced. *Culex pipiens* f *molestus*, was present both within and outside of urban areas in several sites in southern Sweden. In one site, hybrids between the two bioforms were found. *Culex pipiens* f *molestus* has previously been found in urban areas in Sweden, but the detection of the bioform in several rural areas was surprising, indicating that it may be more widely spread than previously thought.

## Introduction

The mosquito species *Culex pipiens* is represented in Europe by two bioforms, *Cx. pipiens* f *pipiens* Linnaeus (L), 1758 and *Cx pipiens* f *molestus* Forskal (F), 1775 (Diptera: *Culicidae*) (Harbach, Dahl et al. 1985) They can act as bridge vectors that transmit emerging arboviruses such as West Nile fever virus, Japanese encephalitis virus, St. Louis encephalitis virus and eastern equine encephalitis virus from birds to susceptible mammals (Crabtree, Savage et al. 1995, Fonseca, Keyghobadi et al. 2004, Hubálek 2008, Ravanini, Huhtamo et al. 2012).

The two bioforms differ in their behavioural and physiological characteristics, which in turn has a major influence on the transmission of arboviral diseases to humans. *Cx. pipiens* f *pipiens*, also known as the northern house mosquito, oviposits its first batch of eggs after a bloodmeal (anautogenous), located in aboveground habitats (epigeous) in rural areas. It undergoes diapause during winter (heterodynamic), mates in open space (eurygamous) and is known to preferably feed on avian blood (ornithophilic). *Cx pipens* f *molestus* does not require a blood meal for oviposition (autogenous), breeds in underground habitats (hypogeous) in urban areas, although, there is evidence of it occurring in aboveground habitats. It is active during winter (homodynamic), does not require large space to mate (stenogamous) and feeds on human blood (anthropophilic) and other vertebrates (Byrne and Nichols 1999, Vinogradova 2000, Vinogradova 2003). In areas of sympatry, *Cx. pipiens* bioforms can interbreed to form hybrids which have an opportunistic behavior in their feeding habits, as they feed on both humans and birds thus increasing the transmission cycle of arboviral diseases (Rudolf, Czajka et al. 2013, Osório, ZÉ-ZÉ et al. 2014). There is evidence that they serve as bridge-vectors between humans and birds from blood-meal analysis (Fonseca, Keyghobadi et al. 2004, Hamer, Kitron et al. 2008). In northern Europe, they have separate but overlapping habitats and the hybridization rate between *Cx pipiens* f *pipiens* and Cx *pipiens* f *molestus* is lower than in south Europe where habitats overlap more (Vogels, van de Peppel et al. 2015).

Despite the differences in their ecophysiological characteristics, they cannot be distinguished using morphological methods of identification since they lack clear characteristics that separate them (Byrne and Nichols 1999, Weitzel, Braun et al. 2011). Although the two bioforms belong to the same species, differentiating between them is important to understand their vectorial potential and there is need for their accurate identification using molecular methods (Harbach, Harrison et al. 1984).

The cytochrome C oxidase subunit I (COI) gene has been used to accurately identify a number of mosquito species although some closely related species cannot be identified due to inadequate variation in the COI locus (Lilja, Nylander et al. 2017). Restriction enzyme digest of the COI marker have been used to differentiate between *Cx. pipiens* bioforms and *Cx. torrentium* (Shaikevich 2007). However, the COI difference between *Cx. pipiens* bioforms has later been questioned (Danabalan, Ponsonby et al. 2012). A rapid assay has also been able to differentiate hybrids of *Cx. pipiens* and *Cx. quinquefasciatus* in North America by examining numerous polymorphisms in the second intron of ACE-2 region. (Smith and Fonseca 2004).

A collection of eight microsatellite loci (CQ11, CQ26, CxqGT4, CxqGT6b, CxpGT4, CxpGTP, CxpGT12 and CxpGT46) have been used for the identification of the *Cx. pipiens* complex and their hybrids (Fonseca, Keyghobadi et al. 2004). One microsatellite loci, CQ11 has two distinct sequences which makes it possible to separate the *Cx. pipiens* bioforms (Bahnck and Fonseca 2006). Despite CQ11 microsatellite locus being considered a diagnostic marker for the accurate distinction of the two bioforms and their hybrids (Bahnck and Fonseca 2006, Amraoui, Tijane et al. 2012, Osório, Zé-Zé et al. 2014), it may misidentify *Cx. torrentium* (Danabalan, Ponsonby et al. 2012). Thus, where *Cx. torrentium* is present, analysis of COI complements CQ11. Given that the two bioforms can mate and form hybrids and that the CQ11 marker is not shown to be linked to *Cx. pipiens* f *molestus* phenotype. It is possible that *Cx. pipiens* f *pipiens* populations can carry a “*molestus* form” CQ11 genotype. Nevertheless, the CQ11 marker has given robust results when compared to multilocus methods predicting *pipiens* or *molestus* form (Farajollahi, Fonseca et al. 2011, Gomes, Sousa et al. 2013).

The nuclear gene CPIJ001674 was recently identified as having a high degree of variation between *Cx* species (Kim, Lee et al. 2018). This could be a potential new marker to distinguish the two bioforms of *Cx. pipiens*.

*Cx. pipiens* is found in most parts of Sweden (Lundström, Schäfer et al. 2013). *Cx. pipiens* f *molestus* has been found previously in Gothenburg (Hesson, Schäfer et al. 2016) and has been reported from Stockholm in 1934, reviewed in (Lindström 2017). There are also findings of three *Cx. pipiens* f *molestus* from a farm and two wetlands outside Linköping in central south Sweden and hybrids in Linköping (Vogels, Möhlmann et al. 2016). However, there are no other studies that suggest that *Cx. pipiens* f *molestus* is present outside of urban areas in Sweden.

In this study, we collected *Culex* mosquitoes through citizen science outreach, mainly to people in urban areas experiencing mosquito nuisance. Specimens were also sampled in some areas in the south of Sweden. We determined the specimens as *Cx. pipiens* using COI and distinguished the two bioforms; *Cx. pipiens* f *pipiens* and *Cx. pipiens* f *molestus* using CQ11 microsatellite marker. We further tested whether the CPIJ001674 nuclear DNA marker (Kim, Lee et al. 2018) could distinguish the two bioforms.

## Materials and Methods

### Mosquito collection

Mosquito samples were captured as part of a citizen science outreach (Fånga Myggan) in diverse regions of Sweden where people experiencing mosquito nuisance problems were encouraged to submit specimens. In addition, mosquitoes were collected in several areas in south of Sweden (table 1). From the collections, only mosquitoes morphologically identified as *Culex pipiens* sl. were included in the study. From the samples sent in by the public some samples were in such state that morphological identification was not possible. These specimens were nevertheless included. In total, 111 mosquito samples were analyzed (listed in Supplemental table 1).

### Sample Processing and DNA Preparation

Two to three legs were picked from each mosquito and 30 µL PrepMan ultra reagent solution (Applied Biosystems) was added to each tube. Legs were homogenized with a pellet pestle using a hand-held motorized pestle holder for 30secs. The homogenates were then boiled in a heat block at 100°C for 10 mins, chilled on ice for 2 mins and centrifuged for 3 mins at 15000xg. 20 µL of the supernatant from each tube was transferred to a clean tube marked with name and date.

### DNA amplicon sequencing

A Bio-Rad PCR machine (Applied Biosystems, Thermo Fisher Scientific) was used to amplify the samples. Two primer pairs targeting the mitochondrial COI gene (658bp) were used. The universal primers and HCO2198 (Vrijenhoek 1994). Primers amplifying a longer fragment of the COI gene were also used. Fwd_5_COI (GAAGGAGTTTGATCAGGAATAGT) and Rev_3_COI (TCCAATGCACTAATCTGCCATATTA). CPIJ001674 fragment was targeted for amplification by PCR using primers CPIJ001674_fwd (TGTACGTGGAGCACAAGAGC) and CPIJ001674_rev (TCCGAGTAGACCGAGACCAG). PCR amplification reactions, were carried out as previously described (Lilja, Nylander et al. 2017) shortly reactions were carried out in, 2,5 μM MgCl_2,_ 0,2 μM dNTPs, 0.6 µM primers, 0.75 U AmpliTaq DNA polymerase (Thermo Fisher Scientific), 1 µL of DNA. PCR amplifications were performed, with an initial denaturation at 95°C for 3 mins followed by 39 cycles of denaturization at 95°C for 30 secs, annealing 55°C for 1 min and extension at 72°C for 1 min. Final extension was then performed at 72°C for 5 mins. Amplified PCR products were treated using Exonuclease I (thermo fisher scientific) Fast Alkaline phosphatase (Fermentas life sciences). Purified products were Sanger sequenced by Macrogen, South Korea, Inc in both directions.

CQ11 assay was performed as described previously (Bahnck and Fonseca 2006) and run on 1,2% agarose gel to identify bioforms and hybrids but products were also sequenced to further analyze differences.

### Data Analysis

DNA sequences were edited using Bioedit software, analyzed in MEGA version 6.06 after aligning them with ClustalW (Tamura, Stecher et al. 2013).

Obtained COI sequences were individually identified in the Barcode of Life Data (BOLD) System (Ratnasingham and Hebert 2007). A phylogenetic tree of COI was also created by using the Maximum Likelihood method based on the Tamura 3-parameter model. The analysis involved 79 nucleotide sequences. There were a total of 482 positions in the final dataset. Evolutionary analyses were conducted in MEGA6. Genetic signatures targeted for the COI fragment described in Shaikevich (Shaikevich 2007) were assayed to differentiate between bioforms *pipiens* and *molestus*. CPIJ001674 gene sequences were aligned and compared (Kim, Lee et al. 2018) while CQ11 microsatellite sequences were aligned and compared with publicly available sequences DQ470141, AF075420, DQ470149, EU345579, EU345580, KP675975, KP675979-KP675982.

## Results

### Taxonomy of submitted specimens

In this study, we asked members of the public experiencing problems with mosquitos in urban areas to submit specimens. In addition to this, mosquitoes were collected in rural control areas in southern Sweden. In total, 111 specimens were analyzed. Of the specimens, 58 were classified as *Cx. pipiens* f *molestus*, 3 were hybrids, 32 were *Cx. pipiens* f *pipiens*, 12 were *Cx. torrentium*, 2 were not *Culex* mosquitoes and 4 failed to be classified. Most of the *Culex pipiens* f *molestus* specimens were collected in Gothenburg and the bioform was more spread throughout the city than previously reported and was also present in neighbouring suburban locations. The molestus bioform was also present in the northern Malmoe suburb Burlov and in rural locations such as Sollebrunn and Horby. Hybrids were found in Simrishamn (Fig 1).

**Figure 1).**
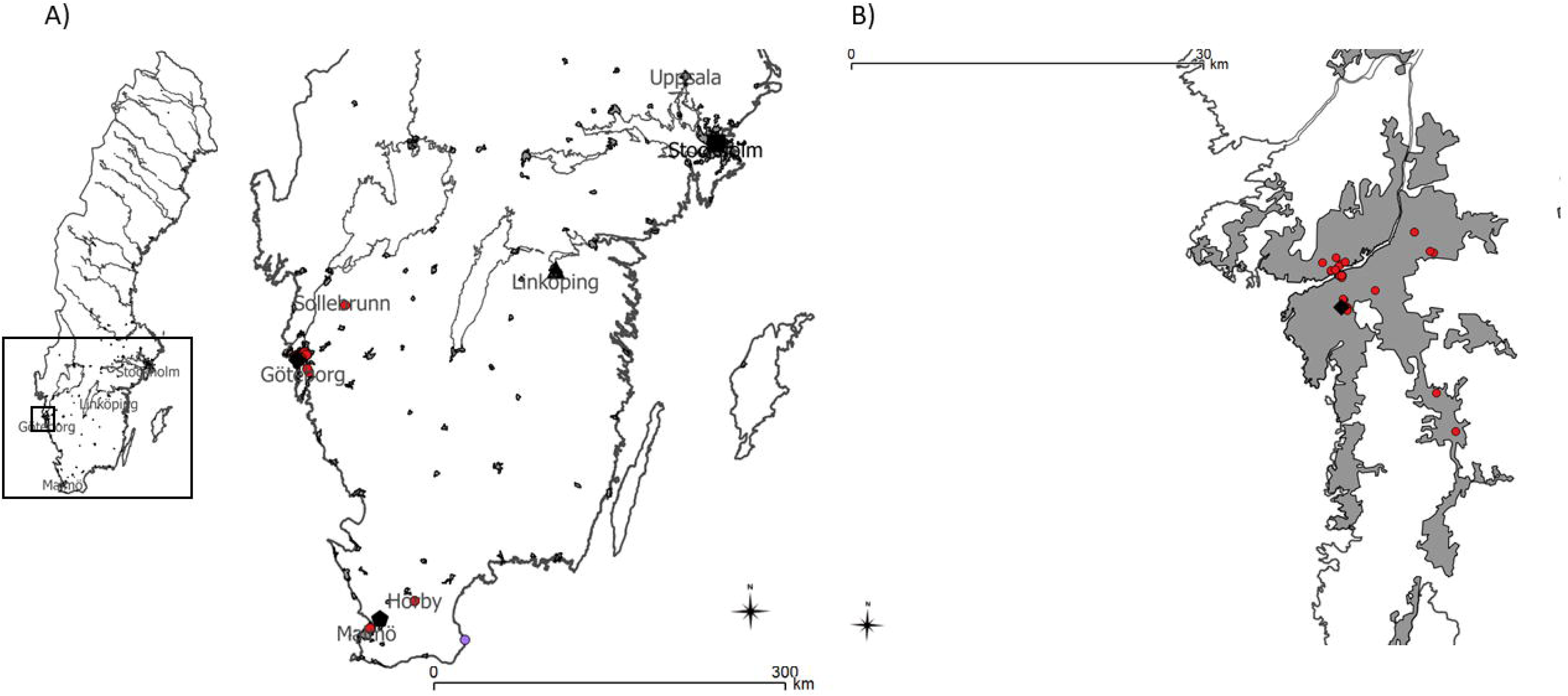
A) Map showing the locations where *Culex pipiens* f *molestus* was found marked with red. Former reports of *Culex pipiens* f *molestus* are marked with black. Square, Stockholm. Forsslund (1941), pentagon, Lund. Rydberg (1933), triangle, Linköping. Vogels (2016) and diamond, Gothenburg. Hesson (2016). B) The spread of *Culex pipiens* f *molestus* in Gothenburg is shown in higher magnification.

### Species identification using the COI gene

Specimens were classified to species level by comparing the COI sequence to published sequences in the Barcode of life database (BOLD.org). COI sequences were also aligned and a phylogenetic tree analysis confirmed the classification into Culex torrentium and Culex pipiens (Fig 2). No specimens had the COI variant suggested by Shaikevich (2007) as diagnostic for *Cx. pipiens* f *molestus* for Russian *Cx. pipiens* mosquitoes, suggesting that this marker was a poor predictor of *Cx. pipiens* f *molestus* in Sweden. Shaikevich also suggested that this site was differing *Cx. torrentium* from *Cx. pipiens* f *pipiens* but interestingly, one specimen of *Cx. torrentium* shared the same variant as that of *Cx. pipiens* f *pipiens*. Nevertheless, there were 12 specimens classified as *Cx.torrentium*.

**Figure 2).**
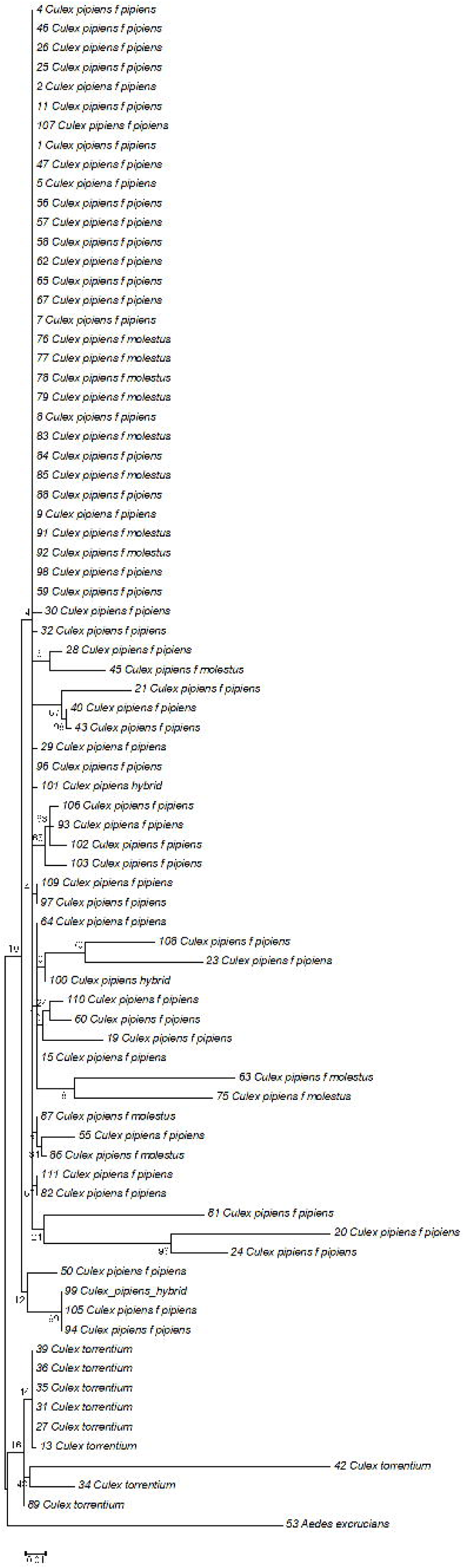
Phylogenetic tree of COI sequences. *Culex pipiens* form a separate cluster from *Culex torrentium*. The phylogenetic tree was inferred using the Maximum Likelihood method based on the Tamura 3-parameter model. The percentage of trees in which the associated taxa clustered together is shown next to the branches. The tree is drawn to scale, with branch lengths measured in the number of substitutions per site. Evolutionary analyses were conducted in MEGA6.

### Bioform classification using the CQ11 microsatellite marker

Analysis of the amplification product of the CQ11 locus was analyzed on agarose gel and each specimen was classified into bioform *pipiens* or *molestus* according to the size of the amplicon. Further the amplicons were sequenced and the sequences from the mosquitoes were compared with publicly available sequences in GenBank and there was a clear distinction between the two *Cx. pipiens* bioforms. Of all the mosquitoes sequenced (n=111), thirty specimens (27%) were classified as *Cx. pipiens* f *pipiens* while fifty-nine specimens (53%) were classified as *Cx. pipiens* f *molestus*. three sequences (3%) were from *Cx. torrentium* and fifteen specimens failed to produce sequence (14%).

### Potential bioform classification using the CPIJ001674 gene

The CPIJ001674 gene was sequenced from 64 mosquito specimens to evaluate this marker as a potential new marker for the *molestus* bioform. Several sites were polymorphic but there was no distinction between *Cx. pipiens* f *pipiens* and *Cx. pipiens* f *molestus*. A phylogenetic tree of the sequences had low bootstrap values and showed no stable clusters (Supplemental fig 1). Whilst the CPIJ001674 gene was published as a gene that showed polymorphisms among the *Culex* complex species mosquitoes, there was no clear distinction in our hands, even between *Cx. pipiens* and *Cx. torrentium* mosquitoes.

## Discussion

*Cx. pipiens* f *molestus*, is a known vector for several pathogens, and can be an efficient bridge vector for flaviviruses such as West Nile fever virus and Usutu virus due to its wide range of hosts. As such, it is of interest to distinguish *Cx. pipiens* f *molestus* from *Cx. pipiens* f *pipiens* which prefers avian hosts over mammals.

In southern Europe, the division between the two bioforms is less pronounced where they share many ecological niches. In northern Europe, *Cx. pipiens* bioforms have been thought to be more isolated from each other and hybrids are less common (Osório, Zé-Zé et al. 2014, Vogels, van de Peppel et al. 2015, Vogels, Möhlmann et al. 2016). Compared to the few previous reports of *Culex pipiens* f *molestus* in Sweden we have found specimens in several rural areas where we would not have expected it to be present. Our study analyzed specimens sent in by people experiencing mosquito nuisance in the south of Sweden and compared with several collections of *Culex* mosquitoes collected in traps in rural areas in the south of Sweden. Most of the *Cx pipiens* f *molestus* specimens were collected in Gothenburg where the bioform has been recently described (Hesson, Schäfer et al. 2016). This makes it hard to evaluate the true distribution of the *molestus* bioform. However, it is clear that it is more common than prevously thought. This indicates that the bioform may be present in even more places if it is investigated more closely. From the mosquitoes sent in by people experiencing nuisance most mosquitoes were *Cx. pipiens* f *molestus* while none of the specimens collected in traps were bioform *molestus*. Contrasting with the results from (Vogels, Möhlmann et al. 2016) where they found the *Cx. pipiens* f *molestus* in rural sites and hybrids in an urban site where no nuisance problems have been reported. Further studies will be needed to more closely study the distribution of *Cx. pipiens* f *molestus* in Sweden.

In this study we sequenced COI to identify the mosquitoes. The COI locus was able to identify *Cx. pipiens* and *Cx. torrentium* specimens. However, it was not able to separate the two *Cx. pipiens* bioforms from each other as suggested by (Shaikevich 2007). The *Cx. pipiens* mosquitoes sequenced in our study all shared a single nucleotide variant in the COI gene which identified Russian *Cx. pipiens* f *pipiens* in the study by Shaikevich. The *Cx. pipiens* f *molestus* in our study did not share the SNP in the COI locus found in Russian *Cx. pipiens* f *molestus*. This was similar to what was previously observed in the UK and US (Kothera, Godsey et al. 2010, Danabalan, Ponsonby et al. 2012). The nuclear DNA marker (CPIJ001674) was not reliable for the distinction of the *Cx. pipiens* biotypes in our study. The gene region was previously suggested to have a nucleotide variation that could be used for differentiating between the bioforms (Kim, Lee et al. 2018) but this study only compared lab strains of *Cx. pipiens* of both bioforms and differences between these might not be reflected in wild caught mosquitoes.

## Conclusions

*Cx. pipiens* f *molestus* is present in several urban and rural locations in Sweden. Further studies of *Cx. pipiens* in Sweden could further clarify the spatial distribution and how this might affect future transmission of mosquito borne viruses such as West-Nile virus and Usutu virus that are spreading in Europe.

## Acknowledgement

The research was co-funded by the Swedish Research Council grant 2017-05607 and the Swedish Civil Contingencies Agency (MSB) that supported the collection and identification of mosquito samples.

## Figure legends

Table 1).

A list of the locations where specimens in the study were collected.

