## Supplementary figures and images for "Human biting *Culex pipiens* bioform *molestus* is spread in several areas in south Sweden"

### Supplemental fig 1

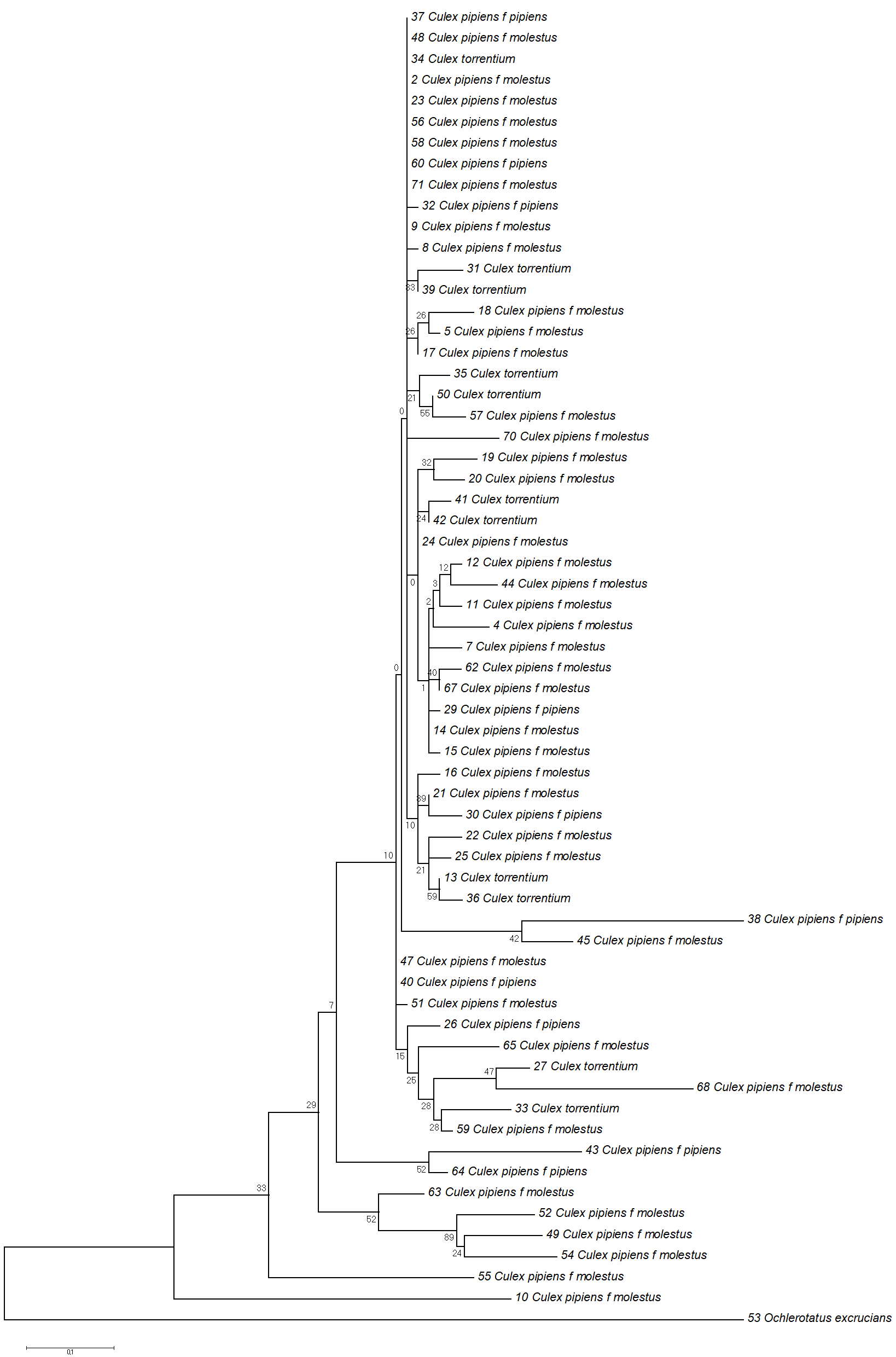
